# Rtrack: a software package for reproducible automated water maze analysis

**DOI:** 10.1101/2020.02.27.967372

**Authors:** Rupert W Overall, Sara Zocher, Alexander Garthe, Gerd Kempermann

**Affiliations:** CRTD – Center for Regenerative Therapies Dresden, Technische Universität Dresden, 01307 Dresden, Germany; DZNE, German Center for Neurodegenerative Diseases, 01307 Dresden, Germany

## Abstract

*Rtrack* is an open-source software package for the analysis of spatial exploration data from behavioural tests such as the Morris water maze. The software provides an easy-to-use interface for data import, analysis and visualisation. A parameter-free machine learning model allows rapid and reproducible classification of spatial search strategies. We also propose a standard export format to enable cross-platform data sharing for the first time in the field.

---

Since its introduction, the Morris water maze task^1^ has become the de facto standard for testing hippocampal function in laboratory rodents. The original task measured the time taken to find a hidden platform or the time spent searching in the vicinity of the platform. Hippocampally-lesioned rats demonstrated a particular deficit in re-learning the task after a platform position change.^2^ The simple measures of goal-finding latency during learning and re-learning trials have since been improved upon by the discovery that certain search patterns are associated with more subtle hippocampal deficits.^3–7^ Swim paths can thus be classified into different strategies reflecting the type of learning being employed.

Automated classification of tracking data (typically collected via image analysis of video recordings) has enabled more complex definitions of learning strategies and several algorithms have been developed over the years.^5,8–12^ However, restrictions on the available software tools, such as accessibility (e.g. expensive and/or difficult to use), compatibility (e.g. only for one computer platform), or reproducibility (e.g. many parameters to be hand-tuned), have resulted in a hesitant uptake by the experimental behaviour community. This led us to re-think the approach and develop an open-source, cross-platform, user-friendly and reproducible solution. Our software, *Rtrack*, is written for the widely-used data analysis R/BioConductor^13^ platform and is freely available from the CRAN repository (https://cran.r-project.org/package=Rtrack) or from https://rupertoverall.net/Rtrack where detailed documentation can also be found.

*Rtrack* accepts coordinate data such as, but not limited to, that exported by a range of video tracking platforms. Several popular formats are currently supported and new formats are being added. In a standard workflow, the coordinate data and arena descriptions are read in and automatically normalised. To aid manageability, all experimental details can also be provided in a spreadsheet to allow batch processing of an entire experiment. Then a range of metrics are calculated (details of the metrics are provided in the package documentation and in Supplementary Data 1) and all data and results are collated into a single R language object. These metrics can be analysed using several inbuilt tools and are used to assign tracks to pre-defined search strategies. Strategy calling is fast, and inbuilt parallelisation tools can make this even faster, allowing large data sets to be easily analysed on standard computer hardware. On our test system (4.2 GHz Intel Core i7) processing 1000 tracks took 45 s using a single-threaded mode (1 CPU core) or 15 s running in parallel using 8 CPU threads. Importantly, the strategy calling is also parameter-free, so that no user optimisation is required and results will always be reproducible from the same data. Additional customised analyses can readily be performed by the researcher as all data are available in the workspace as R language objects. The processed metrics can also be exported to a spreadsheet for analysis with other tools if desired.

Because the *Rtrack* package has been written in R, it benefits from the cross-platform support, availability of additional tools (including parallel processing and advanced statistics) and the powerful graphics provided by the R language. Intuitive R development environments are also available along with many learning resources making use of the package accessible even for users who may not have experience with the R platform. The package itself has been written with the non-programmer in mind and a comprehensive introductory tutorial leads new users through the main functions with detailed examples that can easily be adapted to the user’s own data. A range of publication-ready plots can be easily generated, such as individual paths (Fig. 1a) for quality control purposes, density plots (Fig. 1b) to visualise several paths at once and line graphs to show individual variables such as the inbuilt metrics (Fig. 1c).

**Figure 1.**
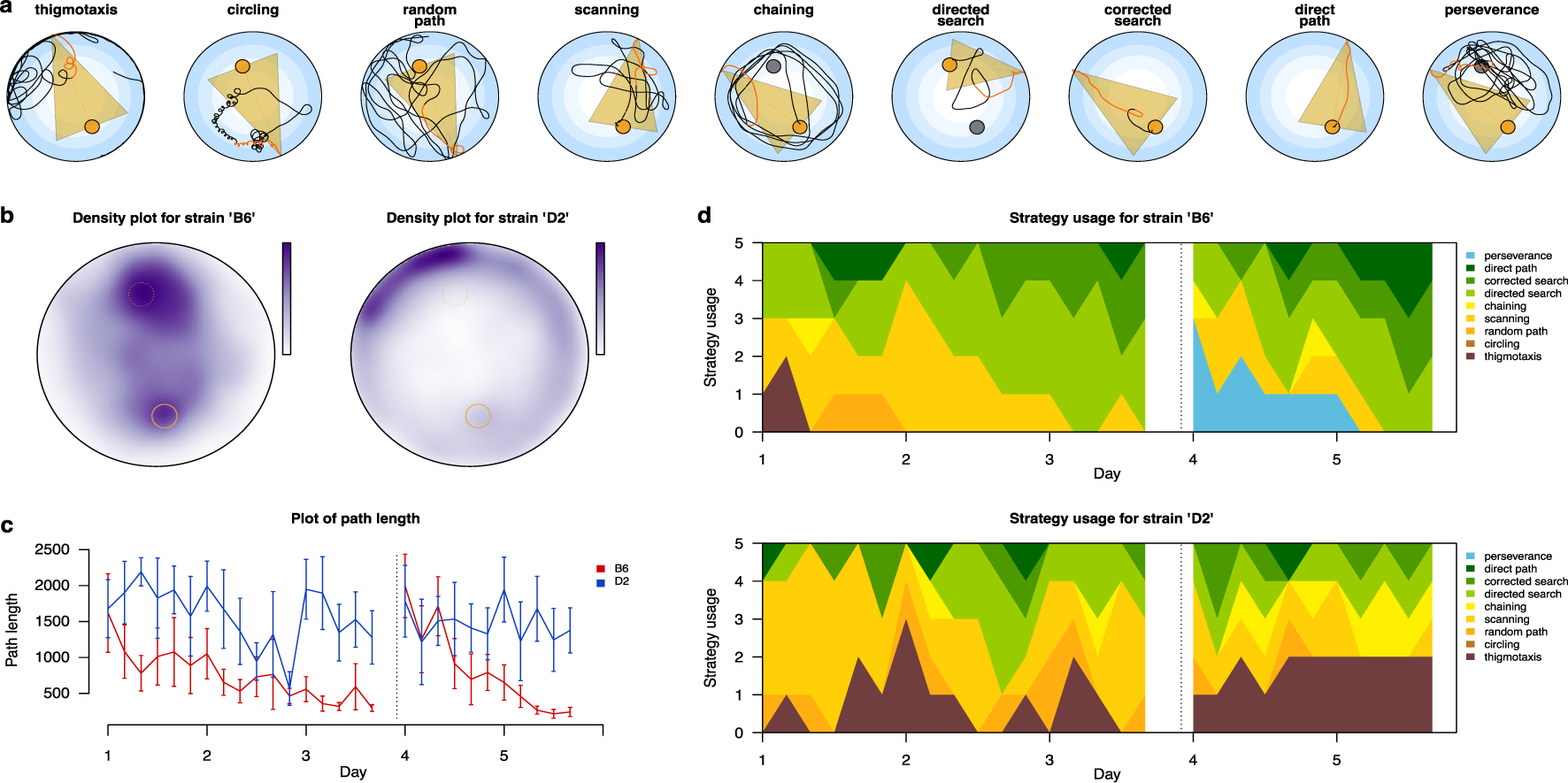
Example plots produced by Rtrack. **a**: Paths representing the nine search strategies. These paths are from the training data set and are those selected by the Rtrack classifier as having the highest confidence for each strategy call. **b**: An example water maze data set with two groups (mouse strains ‘B6’ and ‘D2’) was analysed. Density plots show merged data for all tracks for each mouse strain. Darker shades indicate a higher likelihood of finding the subject in that position. n = 150 tracks for each strain. **c**: Plot of a variable (search path length) where the data have been split by mouse strain. Values are shown as mean ± SEM. n = 5 animals per strain. **d**: Plots of the strategy classification, also split by mouse strain. For these plots, one image is created for each level of the factor (one per strain). Colour boundaries represent cumulative counts of animals using each strategy as indicated on the y-axis. Data from 5 animals are shown in each strain plot. In **c** and **d**, the acquisition (learning) and reversal (change of goal position) phases of the experiment have been separated by a vertical broken line.

Included in the available analysis tools are methods to calculate search strategies. The default approach provided by the package is an automated machine learning classifier, which requires no parameters to be set by the user. To create this classifer, we trained a random forest model using 3111 hand-called strategies from experimental data. This trained model is embedded in the R package and can be used directly to assign processed coordinate data to one of nine spatial search strategies (Fig. 1a). The use of different strategies as the subjects learn the spatial tasks can be monitored and plotted (Fig. 1d) to identify differences between experimental groups that may not be apparent from standard metrics such as latency to goal. A description of the strategies is provided in the package documentation online and in Supplementary Data 2.

The classifier model is able to predict strategies well (Fig. 2a) and calls are typically clear-cut, as evidenced by high confidence compared to the second-choice strategy (Fig. 2b). An indepedent dataset, not used in training the model provided by *Rtrack*, was used to test the machine learning caller (see Online Methods; Supplementary Data 3). A gold standard set of calls was prepared by two experts (authors RWO and SZ) repeatedly manually calling the 280 tracks in this dataset (Supplementary Data 4). These calls were then compared to the calls made by *Rtrack* (Fig. 2c) showing good precision (0.73) and recall (0.66). In fact, the results from the *Rtrack* classifier were no more different to the gold standard than were the calls from each of the two experts from each other (Fig. 2d). This latter point highlights the necessity of a consistent calling algorithm, such as provided by *Rtrack*, in order to provide comparability between laboratories and consistency across large experiments (where it may be unfeasible to rely on a single human caller).

**Figure 2.**
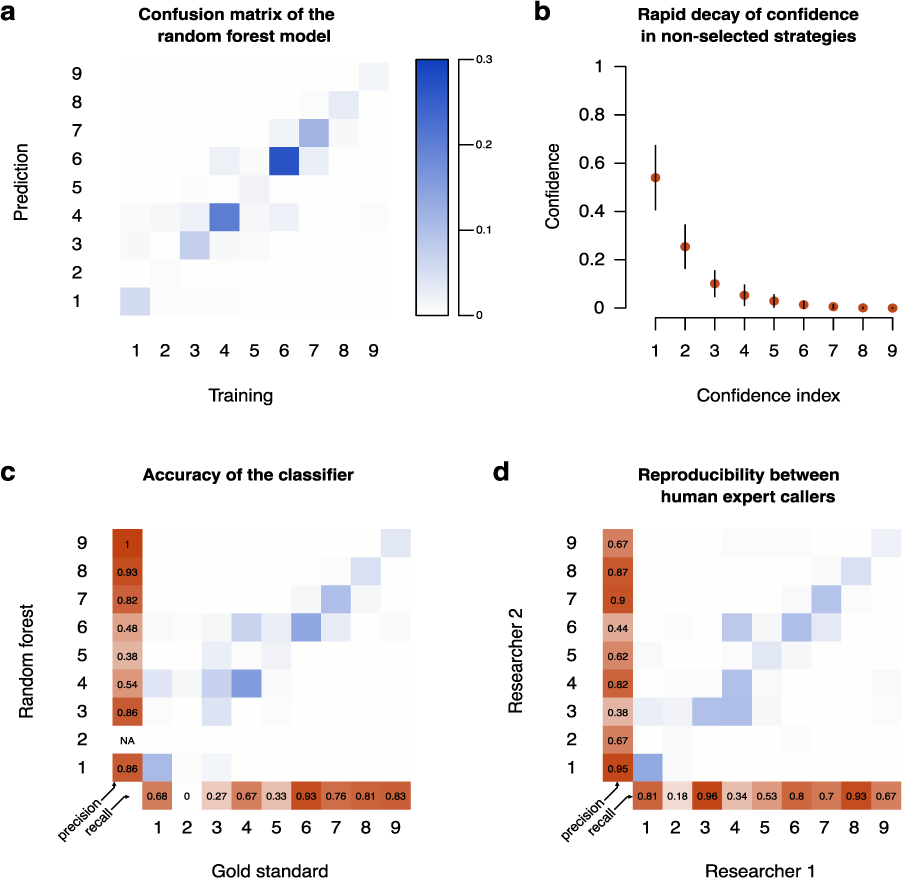
Characterisation of the Rtrack random forest classifier. **a**: Confusion matrix for the model embedded in the Rtrack package. The colour of the grid cells indicates the proportion of all tracks used for training (3111) present in that cell. **b**: Each call was defined as the strategy with the highest confidence score. The confidence of the next best strategy (confidence index = 2) was typically much lower with the confidence rapidly decaying toward zero for other possible strategies. Data are shown as the mean confidence score ± standard deviation. **c**: Strategy predictions were made for an independent dataset for which a ‘gold standard’ had been called by two experts. The random forest classifier built into Rtrack yielded good precision and recall from these data compared to the human-called gold standard. **d**: Calls from the two expert callers were compared showing that differences between human callers were comparable to, or slightly larger than, the difference between the Rtrack model and the gold standard. The grids in panels **c** and **d** use the same normalised colour scale as panel **a** but were generated from a smaller dataset of 280 tracks. For the contingency plots, strategy codes have been used as follows: 1 = thigmotaxis, 2 = circling, 3 = random path, 4 = scanning, 5 = chaining, 6 = directed search, 7 = corrected search, 8 = direct path, 9 = perseverance.

Although the random forest classifier is provided as the default strategy caller in *Rtrack*, the package has been designed to make it easy to drop in alternative callers if desired. For example, an additional classification algorithm has been included in *Rtrack* to aid compatibility with previous published studies.^10^ It will also be possible to include classifiers and analysis tools optimized for other model organisms and for other tasks beyond water maze analysis. The *Rtrack* package is intended to be an extensible platform capable of supporting multiple species and behavioural tasks. Additional modules to support other types of tracking data will be included in future releases.

An important feature of the *Rtrack* package is the ability to export to a universally-readable standard format. All raw data are collapsed into a JSON format text file which can be archived, shared and processed by any other software platform as it is a non-binary, non-proprietry open format. All data required for the reconstruction of an experiment are contained in this archive so that this format can be used for storage of primary raw data. It is concievable that vendors of tracking software may directly offer this format as an export option in the future. A standard data sharing format has so far been missing from the field and the sharing of raw data from Morris water maze experiments is not common—indeed, we could not identify any such data available that would allow reproduction of published analyses. The widespread sharing of data, made possible by the *Rtrack* standard, could enable the integrative re-analysis of behavioural data as has been the case for gene expression since raw data repositories became established.

The *Rtrack* platform enables behavioural researchers with little or no experience in R or programming to rapidly organise their raw data, perform standard and advanced analyses and create publication quality graphics. A data sharing standard for behavioural data is proposed, which will enable re-analysis and comparison of new, and existing, results that has thus far not been possible.

## Online methods

### Calculation of metrics

The *Rtrack* software performs normalisation of raw coordinate data and arena definitions and calculates 34 metrics. The metrics and their calculation are described in detail in Supplementary Data 1 (also available at https://rupertoverall.net/Rtrack/articles/Rtrack_metrics_description.html). Briefly, the supplied arena measurements are normalised to a unit radius and raw coordinate data are correspondingly scaled. Time is scaled to the maximum trial length as provided by the user in the arena definition. Regions of the arena and the path itself are converted into spatial polygons using the R package *sp* enabling complex arena definitions to be supported in the future.

### Training the random forest classifier

A total of 3362 tracks were collated from several unpublished experiments from our laboratory. All tracks were from Morris water maze with mice of mixed sex and genotypes trained for 6 trials per day over 3 days (‘acquisition’) in a 2 m diameter pool followed by 2 further days after a platform position change (‘reversal’). Two experts (SZ and RWO) visually classified images of the swim paths into one of 9 search strategy classes (following the scheme described in Supplementary Data 2; also available at https://rupertoverall.net/Rtrack/articles/Rtrack_strategy_description.html). Each track was classified at least twice and images were repeatedly presented until a consensus call was reached. In cases where a consensus could not be reached (some tracks did not fall into any pre-defined strategy class or consisted of a mixture of strategies), the track was discarded and not used to train the model. In total, 8436 classifications were made which yielded a total of 3111 consensus calls. Images were uploaded to a custom database and web interface and presented randomly so that the researchers were blind to the identity of the animal during classification.

The 3111 calls were used to train a random forest model using the R package *randomForest()*. Some defaults were used to avoid missing data points; ‘latency to goal’ was set to 1 (maximum latency) in cases where the goal was not reached or no goal was present (as in the case of a probe trial), standard deviation of the distance to goal was set to 0 where no goal was present, standard deviation of the distance to old goal was set to 0 where no old goal was present (during acquisition or in the case of a probe trial), and the distance from the goal or old goal used in several metrics was set to 2 (diameter of the normalised arena) in cases where a goal/old goal was not present. A total of 500 trees were grown with 9 variables sampled at each split. The randomForest model object was saved and embedded into the *Rtrack* package.

### Water maze

Data from an independent water maze experiment was used to validate the *Rtrack* classifier on data not used in training the model. In this small experiment, 5 10-week-old female mice each of the strains C57BL/6 (B6) and DBA/2 (D2) had been trained for 6 trials a day for 3 days and for a further 2 days after reversal of the goal position (total of 30 trials per animal). The final acquisition trial (trial 18; 6th trial on day 3) and final reversal trial (trial 30; 6th trial on day 5) were ‘probe’ trials performed without a goal. The water maze consisted of a 2 m pool containing tepid water made opaque with a small quantity of titanium dioxide. The swim paths had been tracked using a video camera attached to a computer running Ethovision 3 (Noldus) and exported to CSV text files. All experiments were conducted in accordance with the applicable European and National regulations (Tierschutzgesetz) and approved by the responsible authority (Regierungspräsidium Dresden; Permit Number: 24-9168.11-1/2009-42).

### Validation of the *Rtrack* classifier

The 280 tracks from the validation experiment (excluding the goal-free ‘probe’ trials) were hand scored using the blinded system described above. From a total of 1314 classifications, 280 consensus calls were obtained. The calls were compared to the results from the *Rtrack* model by creating a contingency table (with the R function *table()*). Macro precision scores were calculated from the diagonal of this table divided by the column sums, and macro recall scores were calculated from the diagonal of this table divided by the row sums of the contingency table. The precision and recall values presented in figure 2 are the arithmetic means of these scores. To assess consistency between the two human classifiers, a similar contingency table was analysed. The results of the gold standard classification, including the data split by researcher, can be found in Supplementary Data 4.

## Data availability

The water maze dataset generated for and analysed during the current study has been deposited in JSON format at figshare with the identifier https://doi.org/10.6084/m9.figshare.10248158.v2^14^, and is also available from the package website in raw https://rupertoverall.net/Rtrack/example_data.zip and JSON https://rupertoverall.net/Rtrack/Experiment.json formats.

## Supplementary Data

Supplementary Data 1. Definition of the metrics used in track analysis (PDF). A document describing the metrics derived from the raw path data. These metrics are the values used to train and query the machine learning classifier. This document is also available online at https://rupertoverall.net/Rtrack/articles/Rtrack_metrics_description.html.

Supplementary Data 2. Definition of the search strategies used by the *Rtrack* classifier (PDF). A document describing the search strategies used by the default *Rtrack* classifier. This document is also available online at https://rupertoverall.net/Rtrack/articles/Rtrack_strategy_description.html.

Supplementary Data 3. Example experiment data (JSON file in ZIP archive). Data from an independent water maze experiment (not used for training the *Rtrack* classification model) was used to verify accuracy of the classifier and perform example analyses and plots. The data from this experiment are contained in a JSON archive produced by the *Rtrack export_json()* function. These data are also available from the figshare repository at https://doi.org/10.6084/m9.figshare.10248158.

Supplementary Data 4. Manual calling of strategies for the gold standard (Excel spreadsheet in ZIP archive). 280 tracks from the example experiment data set (Supplementary Data 3) were manually called by two experts (SZ and RWO). Each track was repeatedly called by both experts until a consensus had been reached (the same strategy had been called at least 3 times). The called strategy (‘call’) is presented alongside the number of times it was called (‘score’), the total number of calls made for that track (‘total’) and the percentage of correct calls (‘score’ / ‘total’). This information was also calculated separately for each expert (‘researcher1_*’ and ‘researcher2_*’).

