## Supplementary Data 1 for "Rtrack: a software package for reproducible automated water maze analysis"

### Description of metrics calculated by the Rtrack package

Rupert W Overall

2020-01-27

At the core of water maze analysis is the summary of the raw paths to yield descriptive metrics. Some of these metrics, such as path length and time to goal are easily explained. Others are less clear, are named differently by different authors and/or depend on variable parameters.

This document describes each of the metrics calculated by `calculate_metrics` and the parameters used in their definition.

#### The metrics object

A core functionality of the Rtrack package is the calculation of metrics from the path coordinates. The `rtrack_metrics` object that is returned by `calculate_metrics` contains the results of these calculations which are used for prediction by the machine learning model. This section explains the information contained in the `rtrack_metrics` object and describes the individual metrics in detail.

##### Object metadata

This section contains information that has been passed to the `calculate_metrics` function and is embedded in the `rtrack_metrics` object for convenience.

#### id

This is the track id supplied either to `read_path` or `read_experiment` and should be a unique identifier for the track. This will be used to name track analysis output and to title the plots generated by `plot_path` or `plot_density`.

###### arena

The `rtrack_arena` object associated with this track. See the function `read_arena` for details.

###### area

A list of the areas of each of the arena components. This is used in several of the metrics calculations.

###### path

The `rtrack_path` object associated with this track. See the function `read_path` for details.

##### Core metrics

This section contains all the metrics, including those calculated for all time points (such as distance from goal and heading error).

###### path.length

The length of the path.

**velocity**

The speed of movement at each time interval.

**total.time**

The total time the subject spent in the arena.

**latency.to.goal**

The time at which the subject first crossed the goal zone. This is often not recognised as a true goal (if the subject does not stop at the goal or if the goal definition is not perfectly calibrated) and so may not be the same as the total time in the arena (even though a trial should be terminated on reaching the goal.

**goal.crossings**

Zone crossings are the number of times the path crossed (i.e. entered and exited once) a particular zone. This is measured by calculating the intersection of the path and the zone polygon and identifying the transitions between 'inside' and 'outside' of the zone. Half of these transitions will be entries which is equivalent to the number of crossings (this value is in fact rounded up in case the track ends in a goal zone, as expected for the goal, which should be counted as a 'crossing').

**old.goal.crossings**

The same as `goal.crossings` for the old goal.

**coverage**

The fraction of the arena covered by the path. This is measured by calculating the area of the minimal polygon enclosing the path (i.e. the 'silhouette') as a fraction of the area of the arena.

**outliers**

The percentage of the path that is not in the not in approach corridor.

**initial.path**

The initial path is the section of the path from the start to a length equal to the direct distance between the start and the centre of the goal.

**initial.heading.error**

The angle between the direct path from start to goal and each section of the initial path.

**initial.displacement.error**

The distance between each point of the initial path and the corresponding point on the direct path from start to goal.

**initial.trajectory.error**

The distance between the last point of the initial path and the centre of the goal.

**turning**

This is calculated for each point as the difference between the angle to the next point and the angle to the previous point. Negatively-signed values indicate a left-hand displacement.

##### turning.absolute

This is calculated in the same way as `turning` but uses absolute values and thus does not contain information about the directionality of the turn.

##### efficiency

The percentage of `initial.heading.error` values below 15°.

##### roaming.entropy

This is an application of the Shannon entropy measure to spatial paths. In Rtrack, it is estimated by calculating a density map with a $50 \times 50$ grid (and a bandwidth of 1/50) which is clipped to the bounds of the arena. This gives a computationally fast estimate of $p$; the likelihood of the path passing through any one of the grid cells. The entropy is then calculated using Shannon's equation: $-\sum(p \times log(p))$. This value is then normalised by dividing by the log of the total number of grid cells present in the arena (so that the final value ranges between 0 and 1). A roaming entropy of 1 means that the path covers the entire arena and it is impossible to predict a point on the path. A roaming entropy of 0 occurs when the entire path is within one grid cell. Lower entropy values indicate a more predictable path. The idea behind the roaming entropy measure is demonstrated at http://www.brandmaier.de/roamingentropy/.

##### time.in.zone.pool

The fraction of the path spent in the pool zone. The pool is defined as the circular arena in which the water maze was performed. The raw size, shape and centre point (to allow for path recording calibration) are set by the experimenter in the arena definition file/s (see the function `read_arena`). The fraction of time spent in this zone should be 1 (excepting only small calibration errors).

##### time.in.zone.wall

The fraction of the path spent in the wall zone. The wall is defined as a ring with the outer extent bounded by the pool/arena and the inner radius equal to 80 % of the arena radius (i.e. the wall zone width is 10 % of the pool/arena diameter).

##### time.in.zone.far.wall

The fraction of the path spent in the far wall zone. The far wall is defined as a ring with the outer extent bounded by the wall (see above) and the inner radius bounded by the annulus (see below).

##### time.in.zone.annulus

The fraction of the path spent in the annulus zone. The annulus is defined as a ring, centred on the centre of the pool/arena, with a width such that the entire goal exactly fits inside it. In other words, the inner radius of the annulus is the minimal distance between the edge of the goal zone and the arena centre, and the outer radius is the maximal distance between the edge of the goal zone and the arena centre.

##### time.in.zone.goal

The fraction of the path spent in the goal zone. The goal size and position are defined by the experimenter in the arena definition file/s (see the function `read_arena`).

##### time.in.zone.old.goal

The fraction of the path spent in the old goal zone. The old goal size and position (if present) are defined by the experimenter in the arena definition file/s (see the function `read_arena`).

**time.in.zone.n.quadrant**

The fraction of the path spent in the north quadrant. The north quadrant is the quarter of the arena area that is centred on the goal. The other quadrants are defined based on this.

**time.in.zone.e.quadrant**

The fraction of the path spent in the east quadrant. The east quadrant is a quarter of the arena area defined relative to the north quadrant (see above).

**time.in.zone.s.quadrant**

The fraction of the path spent in the south quadrant. The south quadrant is a quarter of the arena area defined relative to the north quadrant (see above).

**time.in.zone.w.quadrant**

The fraction of the path spent in the west quadrant. The west quadrant is a quarter of the arena area defined relative to the north quadrant (see above).

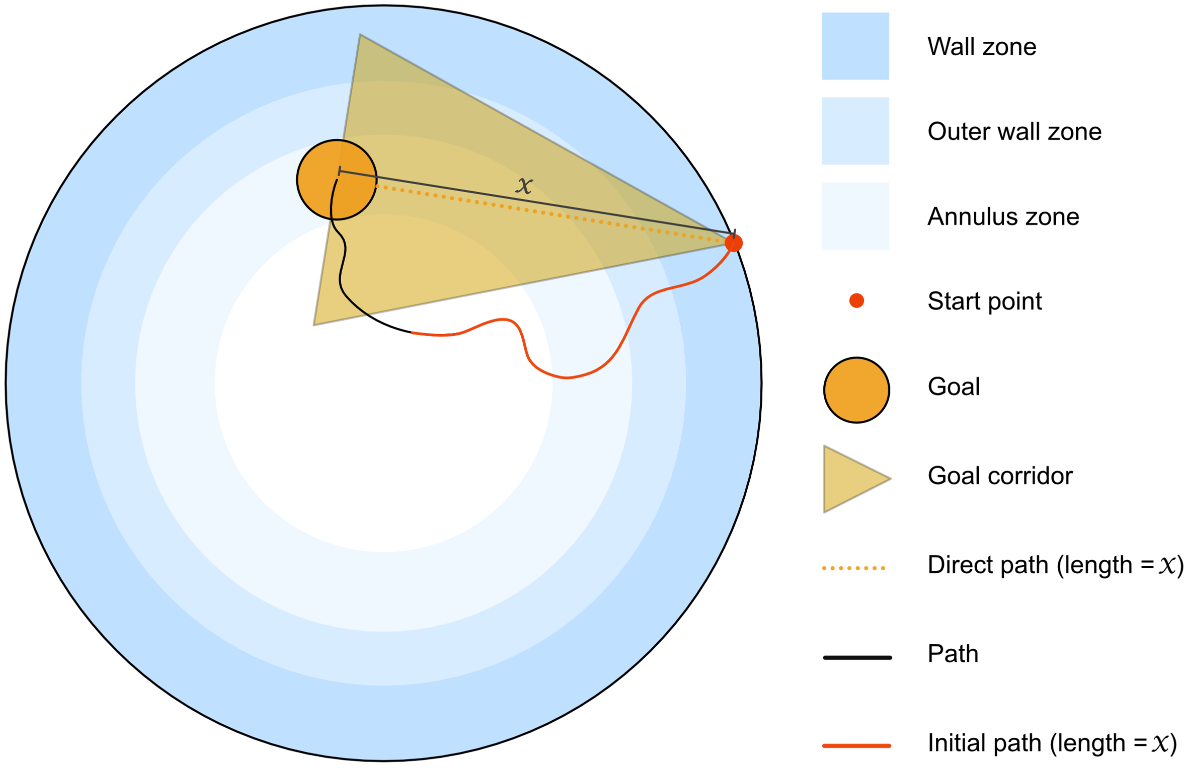


#### Summary metrics

The summary metrics are derived from the core metrics above, but are each only a single value (typically the mean).

**summary**

This vector of values is used for strategy calling using the machine learning model implemented in `call_strategy`.

**mean.velocity**

The mean `velocity`.

**total.time**

The total time the subject spent in the arena (see above).

**latency.to.goal**

The time at which the subject first crossed the goal zone (see above).

**goal.crossings**

The number of times the path crossed the goal zone (see above).

**old.goal.crossings**

The number of times the path crossed the old goal zone (see above).

**coverage**

The fraction of the arena covered by the path (see above).

**mean.d.centroid**

The centroid of the path (the point defined by the mean of all $x$-coordinates and the mean of all $y$-coordinates) is calculated, and then the distance of each point of the path to the centroid calculated. The mean of these values is the `mean.d.centroid`.

**mean.d.goal**

The mean distance of each point on the path from the centre of the goal.

**mean.d.old.goal**

The mean distance of each point on the path from the centre of the old goal.

**mean.d.origin**

The mean distance of each point on the path from the centre of the arena.

**sd.d.centroid**

The standard deviation of the distances of each point on the path from the path centroid (see above).

**sd.d.goal**

The standard deviation of the distances of each point on the path from the goal.

**sd.d.old.goal**

The standard deviation of the distances of each point on the path from the old goal. ##### sd.d.origin The standard deviation of the distances of each point on the path from the centre of the arena.

**centroid.goal.displacement**

The distance between the path centroid and the goal.

**centroid.old.goal.displacement**

The distance between the path centroid and the old goal.

**mean.initial.heading.error**

The mean of the angles between the direct path from start to goal and each section of the initial path (see above).

**initial.trajectory.error**

The distance between the last point of the initial path and the centre of the goal.

**initial.reversal.error**

The distance between the last point of the initial path and the centre of the old goal. This is the same as `initial.trajectory.error` (and uses the same initial path) but records the distance to the old goal (if present). If no old goal has been defined, this value is `NA`.

**turning**

The mean of the `turning` values (see above).

**turning.absolute**

The mean of the `turning.absolute` values (see above).

**efficiency**

The search efficiency (see above).

**Time in zones**

```
time.in.zone.pool
time.in.zone.wall
time.in.zone.far.wall
time.in.zone.annulus
time.in.zone.goal
time.in.zone.old.goal
time.in.zone.n.quadrant
time.in.zone.e.quadrant
time.in.zone.s.quadrant
time.in.zone.w.quadrant
```

Time in zone measures the fraction of a path that is spent in a particular zone. See above for definitions of the zones.

**Unscaled summary metrics**

The same metrics as in `summary`, but many of the values have been returned to the scale of the original data. Because all path and arena coordinates are normalised for use within Rtrack, certain absolute values (such as path length or latency to goal) will not have their original units. This section restores that information and the values plotted by `plot_variable` use these values.

**unscaled.summary**

*See* `summary` *above.*

#### Structure of metrics object

Below is an overview of the hierarchy of the `rtrack_metrics` object together with the names and classes of each component. Where the class is not part of the R base package, it is given in square brackets after the class name.

```
metrics : rtrack_metrics [Rtrack]
    id : character
    arena : rtrack_arena [Rtrack]
        type : character
        description : data.frame
            type : character
            trial.length : character
            arena.bounds : character
            goal : character
            (old.goal : character) **this component is optional and may be missing**
        correction : list
            t : numeric
            x : numeric
            y : numeric
            r : numeric
        pool : list
            x : numeric
            y : numeric
            radius : numeric
            shape : character
        goal : list
            x : numeric
            y : numeric
            radius : numeric
            shape : character
        old.goal : list
            x : numeric
            y : numeric
            radius : numeric
            shape : character
        zones : list
            pool : SpatialPolygons [sp]
            wall : SpatialPolygons [sp]
            far.wall : SpatialPolygons [sp]
            annulus : SpatialPolygons [sp]
            goal : SpatialPolygons [sp]
            old.goal : SpatialPolygons [sp]
            n.quadrant : SpatialPolygons [sp]
            e.quadrant : SpatialPolygons [sp]
            s.quadrant : SpatialPolygons [sp]
            w.quadrant : SpatialPolygons [sp]
            goal.corridor : SpatialPolygons [sp]
    area : list
        pool : numeric
        wall : numeric
```

```
        far.wall : numeric
        annulus : numeric
        goal : numeric
        old.goal : numeric
        n.quadrant : numeric
        e.quadrant : numeric
        s.quadrant : numeric
        w.quadrant : numeric
        goal.corridor : numeric
    path : rtrack_path
        raw.t : numeric
        raw.x : numeric
        raw.y : numeric
        t : numeric
        x : numeric
        y : numeric
        id : character
    path.length : numeric
    velocity : numeric
    total.time : numeric
    latency.to.goal : numeric
    goal.crossings : numeric
    old.goal.crossings : numeric
    coverage : numeric
    outliers : numeric
    initial.path : numeric
    initial.heading.error : numeric
    initial.displacement.error : numeric
    initial.trajectory.error : numeric
    efficiency : numeric
    alpha : numeric
    heading.error : numeric
    time.in.zone : numeric
    roaming.entropy : numeric
    summary : numeric
        path.length : numeric
        mean.velocity : numeric
        sd.velocity : numeric
        latency.to.goal : numeric
        goal.crossings : numeric
        old.goal.crossings : numeric
        coverage : numeric
        mean.d.centroid : numeric
        mean.d.goal : numeric
        mean.d.old.goal : numeric
        mean.d.origin : numeric
        sd.d.centroid : numeric
        sd.d.goal : numeric
        sd.d.old.goal : numeric
        sd.d.origin : numeric
        centroid.goal.displacement : numeric
        centroid.old.goal.displacement : numeric
        mean.initial.heading.error : numeric
```

```
        initial.trajectory.error : numeric
        initial.reversal.error : numeric
        turning : numeric
        turning.absolute : numeric
        efficiency : numeric
        time.in.zone.pool : numeric
        time.in.zone.wall : numeric
        time.in.zone.far.wall : numeric
        time.in.zone.annulus : numeric
        time.in.zone.goal : numeric
        time.in.zone.old.goal : numeric
        time.in.zone.n.quadrant : numeric
        time.in.zone.e.quadrant : numeric
        time.in.zone.s.quadrant : numeric
        time.in.zone.w.quadrant : numeric
        roaming.entropy : numeric
    unscaled.summary : numeric
        path.length : numeric
        mean.velocity : numeric
        sd.velocity : numeric
        latency.to.goal : numeric
        goal.crossings : numeric
        old.goal.crossings : numeric
        coverage : numeric
        mean.d.centroid : numeric
        mean.d.goal : numeric
        mean.d.old.goal : numeric
        mean.d.origin : numeric
        sd.d.centroid : numeric
        sd.d.goal : numeric
        sd.d.old.goal : numeric
        sd.d.origin : numeric
        centroid.goal.displacement : numeric
        centroid.old.goal.displacement : numeric
        mean.initial.heading.error : numeric
        initial.trajectory.error : numeric
        initial.reversal.error : numeric
        turning : numeric
        turning.absolute : numeric
        efficiency : numeric
        time.in.zone.pool : numeric
        time.in.zone.wall : numeric
        time.in.zone.far.wall : numeric
        time.in.zone.annulus : numeric
        time.in.zone.goal : numeric
        time.in.zone.old.goal : numeric
        time.in.zone.n.quadrant : numeric
        time.in.zone.e.quadrant : numeric
        time.in.zone.s.quadrant : numeric
        time.in.zone.w.quadrant : numeric
        roaming.entropy : numeric
```
