## Supplementary Data 2 for "Rtrack: a software package for reproducible automated water maze analysis"

### Description of the strategies called by the Rtrack classifier

Rupert W Overall

2020-01-27

#### Introduction

The core Rtrack classifier provided by this package assigns one of 9 strategies to each path. The strategies are described here together with some notes on their interpretation.

#### Non-goal-oriented strategies

These strategies are not driven by an attempt to find the goal. Typically they are motivated by a desire to escape the testing arena, or by anxious immobility (to the extent that this is possible in a traditional water maze).

##### Strategy 1: Thigmotaxis

Thigmotaxis is an anxiety-related behaviour and is characterised by a swim path that hugs the outer wall or where the subject comes repeatedly in contact with the wall. The subject does not move in a goal-directed manner but rather attempts to get out of the testing arena.

##### Strategy 2: Circling

The subject moves in repeated tight circles. This is distinct from one or two full-turn circles that are sometimes used by the subject to scan the surroundings and reorient. Circling may be anxiety-related.

##### Strategy 3: Random

The subject moves aimlessly throughout the arena with no attempt to search for a goal (or even awareness that a goal exists).

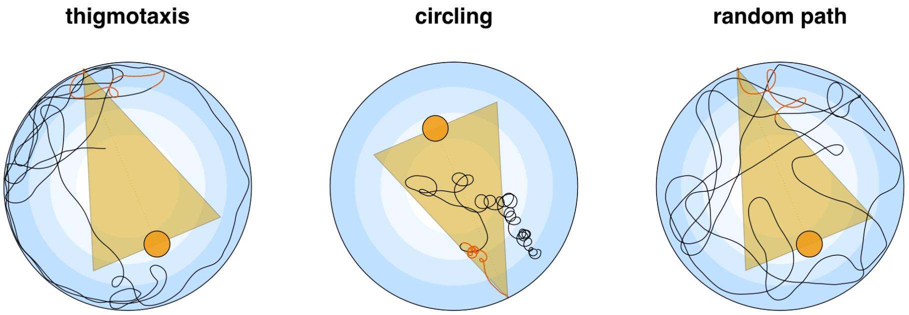


#### Procedural strategies

These strategies involve the use of egocentric behaviours, where the search context is relative to the subject itself. As there should be no local tactile, olfactory or visual information and because the subject should start in different positions relative to the goal, such strategies are unlikely to be consistently successful.

**Strategy 4: Scanning**

The subject moves in repetitive patterns, covering a broad region of the arena. The patterns can either be a 'rolling loop' where the centre of the loop continually changes, or a 'zig-zag'. This behaviour can be localised but off-target or be more goal-focussed.

**Strategy 5: Chaining**

In this pattern, the subject has identified that the goal is located at a specific distance from the wall and thus moves at this distance in the hope of stumbling across it. It can be difficult to detect as the subject will often encounter the goal in less than one full circuit of the arena.

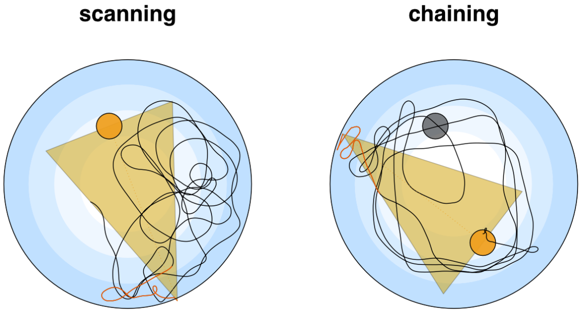


#### Allocentric search strategies

These strategies require the subject to orient themself using distal cues that are in the same frame of reference as the goal. Once a cognitive representation of the distal environment (a 'spatial map') has been constructed, local changes in start position become unimportant and a direct path to the goal can be rapidly calculated. A spatial map also allows a change in goal position to be quickly re-learned.

**Strategy 6: Goal-directed search**

The subject moves in a goal-oriented manner, but not necessarily always correctly. There may be orientation loops, path corrections and path retracing for re-orientation employed. Different from scanning, the path is not repetitive or covering a large area.

**Strategy 7: Corrected search**

The subject moves in an almost direct path to the goal, but makes one or two mistakes, which are corrected by reorientation. Errors can be a small loop to quickly scan the environment to allow re-orientation ('orientation loop'), a course change ('path correction') or a larger loop where the subject returns to a previous point in the path and attempts the search again ('re-orientation bight').

**Strategy 8: Direct path**

The subject moves in a direct line to the goal.

**Strategy 9: Perseverance**

The search pattern involves a repeated re-visiting of the goal location. This strategy is only called when the goal is not present. In the case of a probe trial, where no goal is present, this would be considered a focal search. In the case of a goal position change, this strategy is termed 'perseverence'.

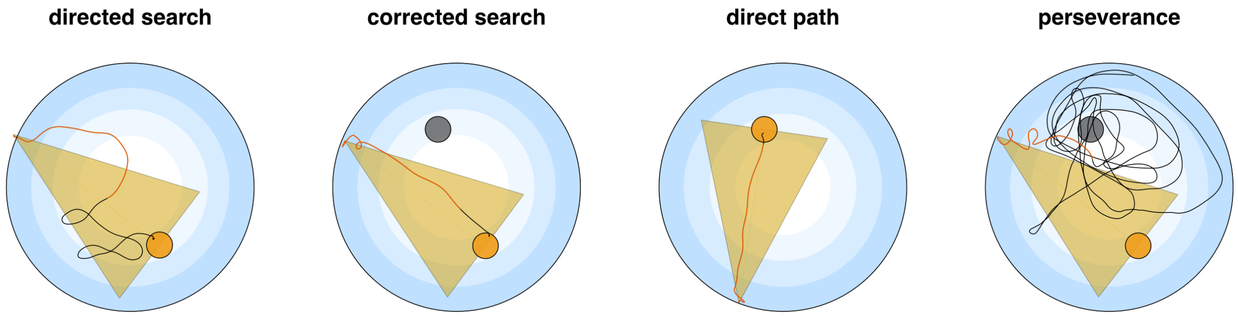
directed search
corrected search
direct path
perseverance
